# Identify and Predict Environmental Change Effects On Tiger Mosquitos, *Aedes Polynesiensis*

**DOI:** 10.1101/284174

**Authors:** Florian Grziwotz, Jakob Friedrich Strauß, Chih-hao Hsieh, Arndt Telschow

**Affiliations:** Institute for Evolution and Biodiversity, Westfalian Wilhelms-University, Münster, Germany, 48149; Institute of Oceanography, National Taiwan University, Taipei, Taiwan, 10617; Institute of Ecology and Evolutionary Biology, National Taiwan University, Taipei, Taiwan, 10617

## Abstract

To control mosquito populations for managing vector-borne diseases, a critical need is to identify and predict their response to causal environmental variables. However, most existing attempts rely on linear approaches based on correlation, which cannot apply in complex, nonlinear natural systems, because correlation is neither a necessary nor sufficient condition for causation. Appling empirical dynamic modelling that acknowledges nonlinear dynamics on nine subpopulations of tiger mosquitos from three neighbouring reef islets of the Raiatea atoll, we identified temperature, precipitation, dew point, air pressure, and mean tide level as causal environmental variables. Interestingly, responses of subpopulations in close proximity (100-500 m) differed with respect to their causal environmental variables and the time delay of effect, highlighting complexity in mosquito-environment causality network. Moreover, we demonstrated how to explore the effects of changing environmental variables on number and strength of mosquito outbreaks, providing a new framework for pest control and disease vector ecology.

## Introduction

Mosquito-borne diseases contribute significantly to human mortality causing more than 1 billion cases and over 1 million deaths every year (GBP 2016). A case in point are mosquitoes of the genus *Aedes* being the main vector of dengue fever. For predicting disease outbreaks, population dynamics of mosquitoes are important concern. A critical need is therefore to understand and predict how environmental change may impact on fluctuation of mosquito populations (Duncombe et al., 2013).

There are two main strains of research aiming to examine how mosquito populations respond to climate changes. On the one hand, lab experiments are conducted to detect causal environmental drivers of mosquito survival, larval developmental time and other fitness parameters (e.g., Galun and Fraenkel, 1961; Couret et al., 2014). However, whether the findings in lab experiments can be generalized to natural systems remain unclear. On the other hand, time series data of mosquito abundance in natural systems are analysed for statistical associations with climate variables. For example, population abundance of the yellow fever mosquito *Ae. aegypti* was found to correlate with temperature, humidity, and precipitation (Scott et al., 2000; Azil et al., 2010; Barrera et al., 2011; Bashar et al., 2014; Alencar et al., 2015, Chaves et al., 2012). However, majority of the studies linking natural mosquito abundances with climate variables employed linear correlation analyses. While, correlation does not necessarily imply causation, lack of correlation does not imply lack of causation either (Sugihara et al., 2012; Chang et al., 2017). Applying linear methods to time series data generated from nonlinear processes bears the risk to detect mirage correlations, which may lead to wrong and misleading conclusions (Sugihara et al., 2012; Deyle et al., 2013; Deyle et al., 2016).

Indeed, mirage correlation (or non-stationary relationship) between mosquito populations and climate variables were often found in nature (Chaves et al., 2012; Simoes et al., 2013). Mirage correlation is commonly found, as a result of a fundamental property of nonlinear dynamical systems, known as state dependency (Sugihara et al 2012; Ye et al., 2015). State dependency means that the relationships among interacting variables change with different states of the dynamical system (Ye et al., 2015). The following example illustrates state-dependence in mosquitoes. It was shown experimentally that *Ae. aegyptii* females have lower vector competence for dengue virus in lower temperature environments only when the temperature is fluctuating; in comparison, without such a specific temperature combination (i.e. mosquitoes living in the environment of high temperature or constant temperature), the females have higher vector competence (Lambrechts et al., 2011; Carrington et al., 2013). For *Aedes* mosquitoes, evidence abounds suggesting that nonlinear population dynamics are common. For example, experimental studies show that larval survival depends on larval density (Barrera et al., 2006), temperature (Couret et al., 2014), and the mosquito’s genomic background and microbiome (Mains et al., 2013) in a complex, nonlinear way. In addition, abundance data of lab and field populations are best described by nonlinear mathematical models (Dye 1984b; Legros et al., 2009; Mains et al., 2013). All these studies suggest that, in general, mosquito-environment interactions are complex, nonlinear, and inter-dependent; as such, linear methods are ill-posed for these questions.

Here, we aim at disentangling the causality network between climate variables and mosquito populations and predict how environmental change may impact on the populations. To overcome the limitation of linear approaches, we use the empirical dynamical modelling (EDM) framework, which is specifically designed for analysing time series data with underlying nonlinear dynamics (Ye et al., 2015; Deyle et al., 2016; Chang et al., 2017). To demonstrate the efficacy of EDM framework for disease vectors, we analysed a unique longitudinal data set of the Polynesian tiger mosquito, *Aedes polynesiensis*, which is an important vector for lymphatic filariasis and dengue fever in the South Pacific (Lambdin et al., 2009; Lau et al., 2016) (see supplementary material for more information on *Ae. polynesiensis*). The data consist of biweekly (14-day interval) mosquito counts over two years at nine sampling points on three motu islands (Mercer et al., 2012; Fig. S1, S2). The motus have similar climate, but differ in vegetation and human activity as well as in larval competition and average body size of adult mosquitoes. This exceptionally high temporal and spatial resolution gives unique opportunities for studying population dynamics of mosquitos. In particular, it allows investigating to what extent the local habitat drives mosquito dynamics and how neighbouring population in a meta-population are interconnected. Our objectives are (1) to identify the critical environmental variables that causally affect mosquito populations using convergent cross mapping (CCM) (Methods), (2) to test for causal interactions between mosquito subpopulations, and (3) to explore scenarios how change in a causal environmental variable may affect mosquito abundance as well as number and strength of mosquito outbreaks.

## Results

### Causality between environment and mosquito abundance

Comparing the results based on CCM versus correlation demonstrates the efficacy of using EDM in our data. Indeed, correlation does not imply causation (Fig. S3), while lack of correlation does not imply lack of causation (Fig. S4). Moreover, correlation can change sign, depending on ecosystem context, known as mirage correlation (Figure S5). These examples from our data highlight the need to employ correct methods for studying causality in complex nonlinear systems.

When analysing the mosquito dynamics for each of the nine sampling sites separately using CCM, we found that temperature, precipitation, dew point, air pressure, and mean tide level are causal variables for the mosquito populations (Fig.1; supplementary Table S2). Remarkably, the forcing environmental variables varied substantially among the nine sampling sites (Fig. 1). For example, we found that sampling site S7 at Toamaro island is affected by four climate variables (temperature, dew point, precipitation, mean tide level), site S8 by only one (air pressure), and site S9 by none. Similarly strong differences between sampling sites were found for the other two islands. However, when repeating the same analysis for each of the three islands (sum of the three sites within the same island) and for the whole meta-population (i.e., sum of the nine sampling sites), we found no climatic effect on mosquito abundance from any variable that we analysed here. The spatial scale dependence is remarkable, given the small sizes of the islands (100-500 m length) and short distance between sites (a maximum distance between two sampling sites of less than 5 km).

**Figure 1.**
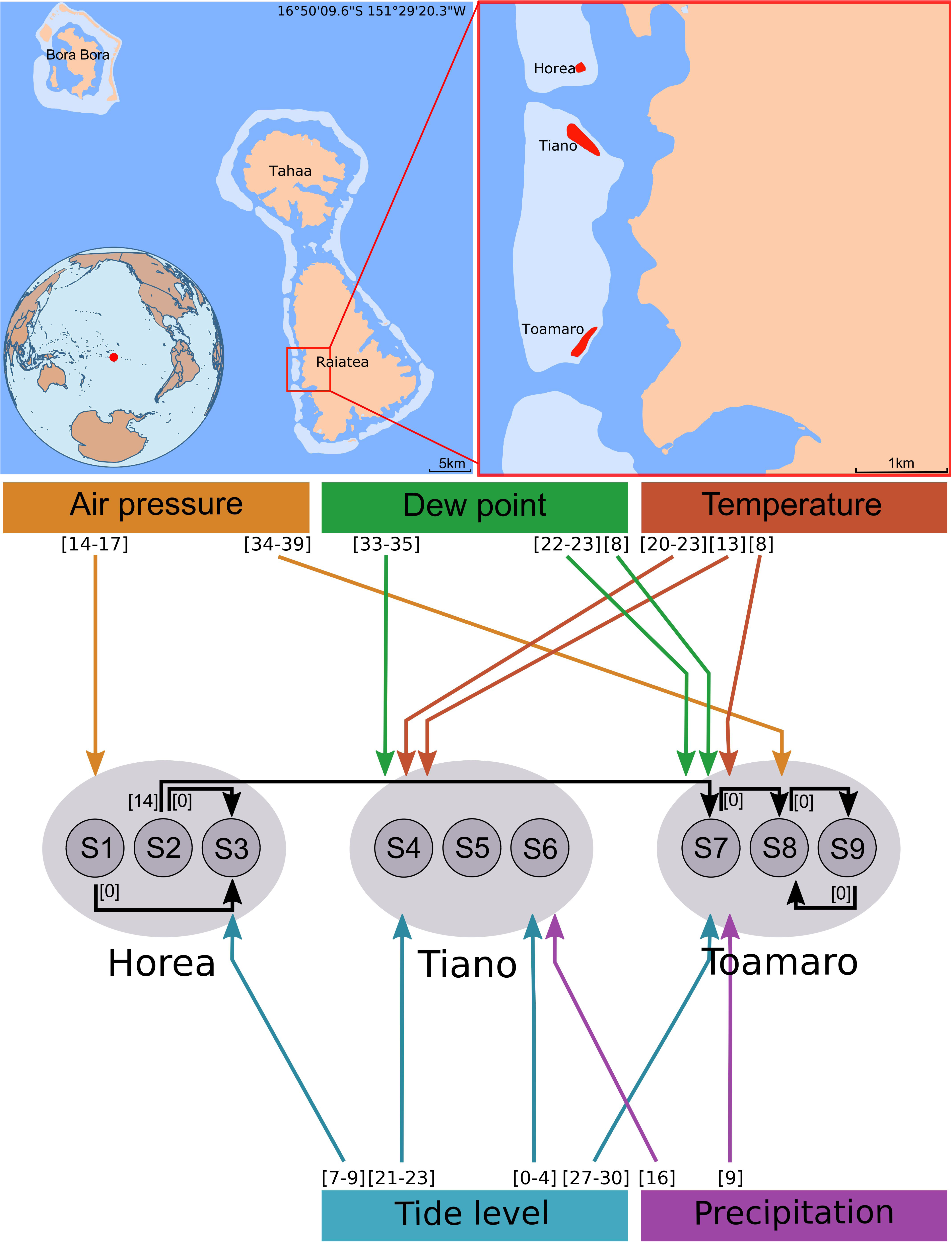
Causality network of the Polynesian tiger mosquito (*Ae. polynesiensis*). *Upper graph:* Field site in Raiatea (French Polynesia). *Lower graph:* Causal interactions within the mosquito meta-population (black arrows) and between climate variables and mosquito abundances (coloured arrows). Causality was evaluated using convergent cross mapping. Only significant results are shown (see Methods, supplementary Tables S2, S3 for details). Numbers next to arrows indicate the time lag *d* (in days), for which causality was detected. Symbols S1 to S9 denote the different sampling sites.

Moreover, the causal forcing variables showed large variation in time lag, from zero day to more than five weeks even for the same environmental variables. For example, the mean tide level affects the mosquito dynamics at S6 with a time lag of 0-4 days, but the lag is 27-30 days at S7; time lags for air pressure are 14-17 days at S1, but 34-39 days at S8. In other words, the causal effect of climate on the mosquito dynamics has strong temporal scale dependence.

A striking spatial pattern emerges when the three islands are compared with respect to the number of causally forcing variables and the time lag of the causal effect. For Horea, we found only the mean tide level and the air pressure to be causal drivers. The associated time lags are 7-9 and 14-17 days, respectively (Fig. 1). For Tiano and Toamaro, on the other side, we detected four and five causally forcing variables, respectively, and the associated time lags range from 0 to 35 days for Tiano and 9 to 39 days for Toamaro (Fig. 1). These results suggest that mosquito-environment interactions are more complex on Tiano and Toamaro than on Horea.

### Causality between mosquito populations

We tested for causal interactions between all pairs of the nine subpopulations, and found bidirectional causality in one case (S8<->S9) and unidirectional causality in four cases (S1->S3, S2->S3, S2->S7, S7->S8) (Fig. 1, black arrows; supplementary Table S3). A clear spatial pattern emerges. All causality links start and end at Horea or Toamaro, but none at Tiano. Further, five of the six links appear within motu (either Horea or Toamaro), with no time lag. Between-motu interaction among subpopulations was only found in one case, from Horea (S2) to Toamaro (S7), with a time lag of 14 days. This case is surprising because Horea and Toamaro are not neighbouring motus, but separated by Tiano (cf. Fig. 1). The lack of connectivity between the subpopulations of Tiano and the other two motus may be explained by the fact that Tiano has the lowest mosquito abundances due to intensive pest control management (Mercer et al. 2012).

### Scenario Exploration

After identifying the critical forcing environmental variables, we then explored the scenarios how changes of these causally forcing variables may affect mosquito abundance (Methods). The key idea is to predict mosquito abundances for scenarios, in which a climate variable is increased or decreased at each time point, and then compare the predicted abundances with the original values. Take site S7 as an example, Figure 2 shows the observed abundances of *Ae. polynesiensis* and the predicted abundances for a scenario with increased mean tide level. We found that the predicted abundances are higher than the observed for 28 time points but lower for the other 19. Note that increasing tide level does not always result in reduced or elevated mosquito abundance; rather, the response depends on the state of population (e.g. the abundance of mosquitos in combination of other environmental variables at a given sampling interval). We conducted a large-scale screen for all causality links between climate variables and mosquito abundance. The results suggest that *state dependence* is a common feature of the *Ae. polynesiensis* population dynamics (supplementary Fig. S7).

**Figure 2.**
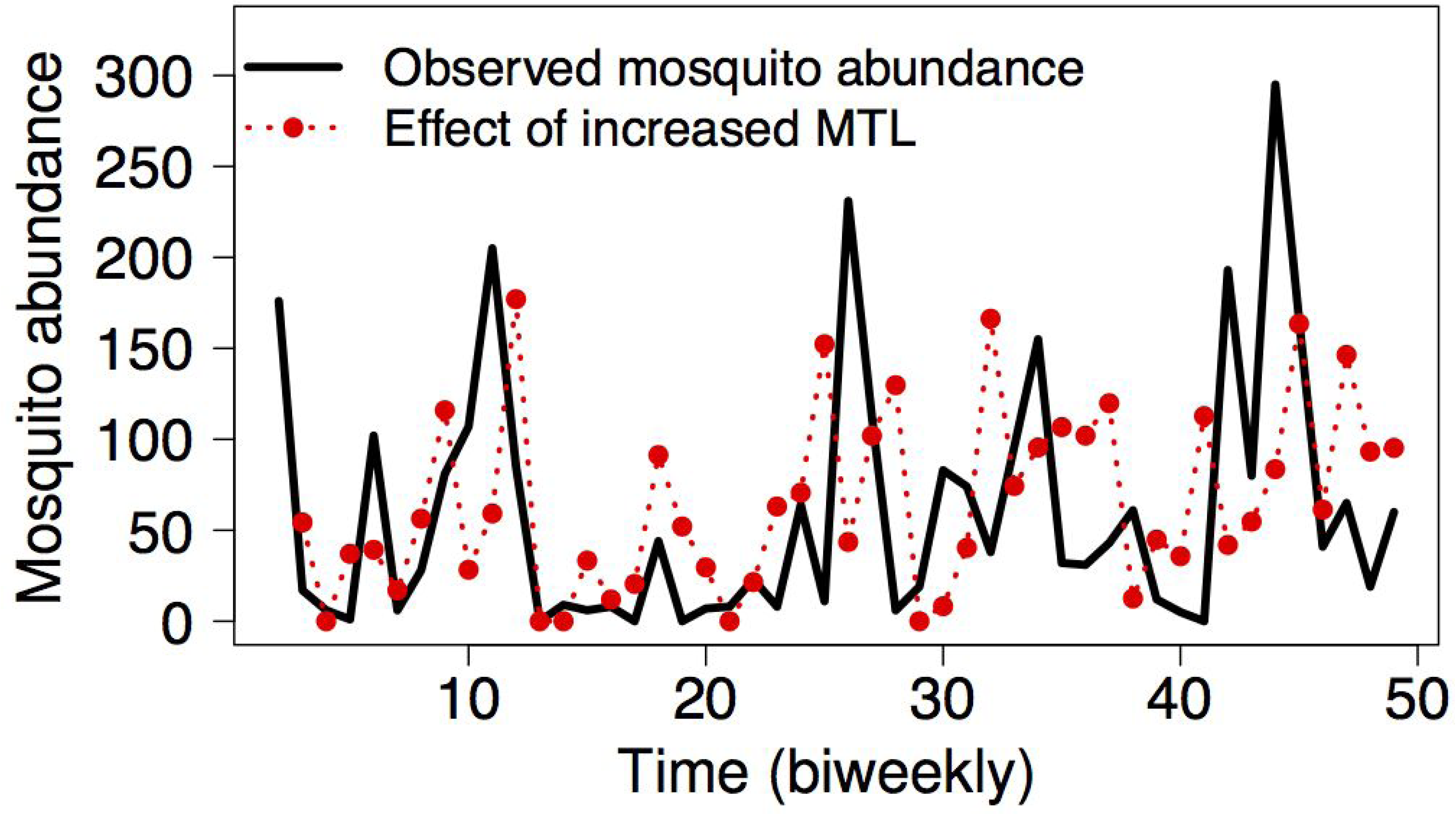
Predicted state-dependent effect of environmental change on mosquito dynamics using scenario exploration. Shown are the observed mosquito abundances at sampling site S7 (black), and the predicted abundances for the same sampling site under an increase of the mean tide level by half of its standard deviation (red). Predictions were generated using multivariate simplex projections and a time lag of *d*=28 days. Note that there is a causal effect of the MTL on the S7 population for this time lag (see Fig. 1). The analysis predicts that an increase in the MTL results in increased or decreased mosquito abundance depending on the state of the system.

Finally, using the same approach, we also explored how environmental change affects the number and strength of mosquito outbreaks. Here, mosquito outbreak is defined as mosquito abundance that is five standard deviations larger than the median abundance of a given time series. This analysis focuses on mosquito peak abundances but not averages because the former are of particular importance for pest control and disease prevention. For example in S6, changes in the mean tide level may have substantial effect on mosquito outbreaks (Fig. 3). Increased mean tide level is predicted to increase the number of mosquito outbreaks from 12 up to 17 (Fig. 3A, C) and the outbreak strength by up to a factor of 1.5 (Fig. 3A, C, D). However, the predicted effect depends on the specific time lag under consideration. For instance, a moderate reduction in MTL is predicted to reduce the number of outbreak for *d*=0 (Fig. 3A), but to increase outbreak for *d*=1 (Fig. 3B). General predictions for the mean tide level are difficult to make because our methodology does not allow evaluating the joined effect of all time lags. At S7, changes in three climate variables were found to significantly alter number and/or strength of mosquito outbreaks (Fig. 3E and F), including the dew point with a lag of 22 days, the mean tide level with a lag of 28 days, and precipitation with a lag of 9 days.

**Figure 3.**
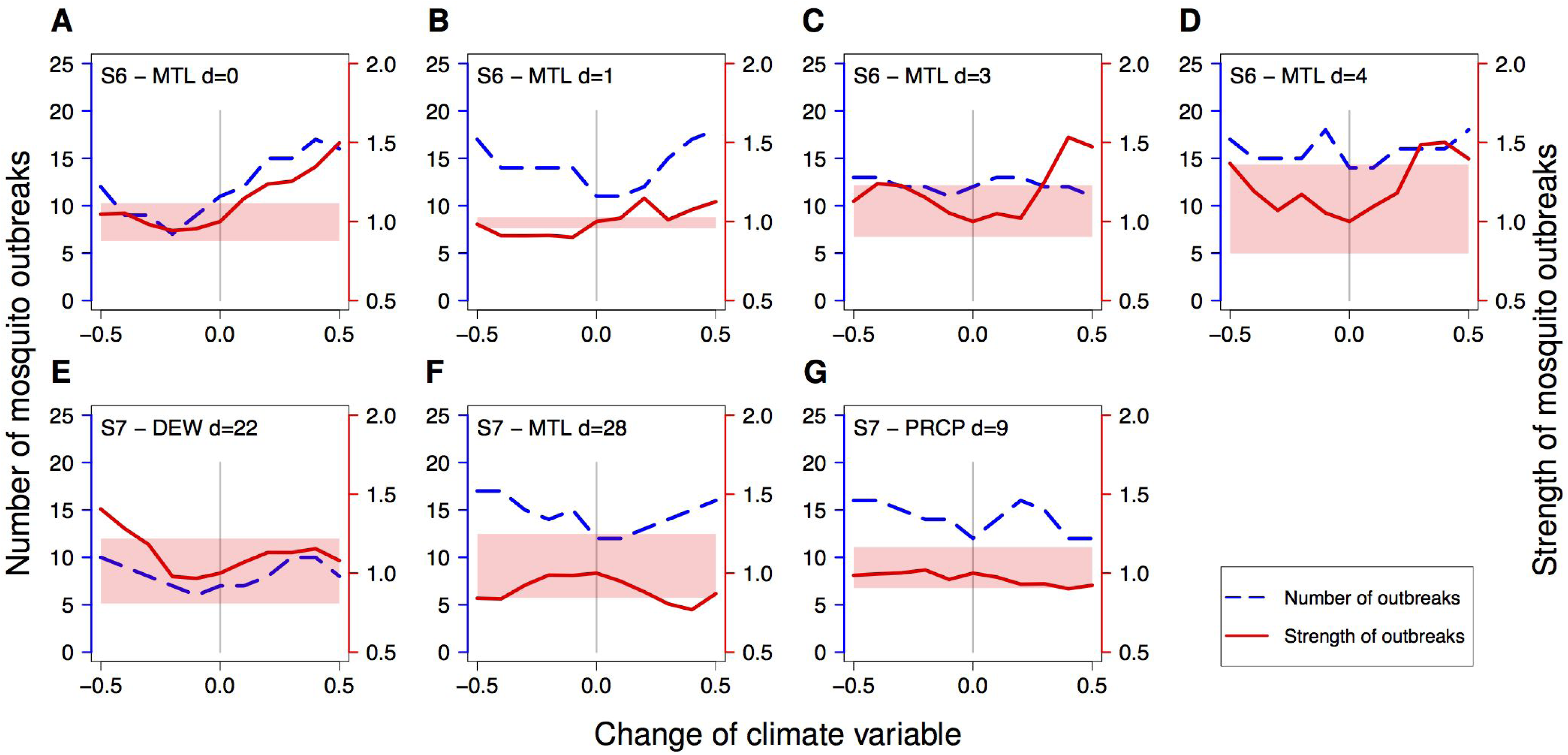
Effect of climate change on number and strength of mosquito outbreaks. Two types of scenario explorations were conducted. The blue line shows how climate change affects the number of mosquito outbreaks, defined as mosquito abundances that are five standard deviations larger than the median of the original data. The red line indicates the relative strength of mosquito outbreaks, defined as the arithmetic mean of the highest five population abundances under climate change divided by that of the original data. The light red zone indicates the range between the highest and the fifth highest mosquito abundance of the original time series. The figure shows only the cases, in which the outbreak strength is at least once outside of the light red zone. These are considered as scenarios with a substantial effect of climate change on mosquito abundance. However, the analysis was conducted for all causality links between climate and mosquito abundance (see supplementary Table S2).

## Discussion

### Causality between environment and mosquito abundance

We applied for the first time the causality test using CCM to a longitudinal mosquito data set and found that temperature, precipitation, dew point, air pressure, and mean tide level are forcing climate variables. Our analyses confirm the common belief that temperature and precipitation causally affect mosquito abundance; in particular, container breeder mosquitoes, such as *Ae. polynesiensis* and *Ae. aegypti*, critically rely on rainfall and moderate temperatures for their larval development (Focks et al., 1993). However, previous rigorous tests for causal effects of environmental factors on mosquitos stem only from lab experiments (e.g., Couret et al., 2014), whereas studies using longitudinal field data employed mostly linear statistical methods such as correlation analysis (Scott et al., 2000; Azil et al., 2010; Alencar et al., 2015) that are ill-posed for the nonlinear mosquito dynamics. Here, our causality analysis confirms the experimental predictions that, however, hitherto have not been confirmed at the population level in the natural environment.

Surprisingly, we found that the mean tide level, an environmental factor that was rarely discussed in the literature, is the most influential environmental variable (quantified by the number of significant tests) in affecting mosquito populations in French Polynesia (Fig. 1). We suggest the effect of tide may be related with the special life style of *Ae. polynesiensis*. Specifically, larvae of *Ae. polynesiensis* can develop in breeding containers filled with brackish water due to their high tolerance to salinity (Ingram 1954; Chang 2010). Considering that tide is likely to affect both the water level and the degree of salinity in the breeding containers, tide potentially has an impact on the number and quality of breeding sites, which may cause changes of oviposition, larval density, larval survival and subsequently mosquito population dynamics. Our finding may have important management implications because the tide level is mainly determined by lunar cycles and therefore bears a high long-term predictability. This is in stark contrast to the other climate variables, which have comparatively low long-term predictability. An interesting question for future research is whether the long-term predictability of the tides may be used for improving forecast of *Ae. polynesiensis* outbreaks.

We additionally found that the air pressure is a forcing variable of *Ae. polynesiensis*. While the effect of air pressure on mosquito survival has been suggested from lab experiments, although for a different species (*Ae. aegypti*; Galun and Fraenkel, 1961), the empirical evidence for natural mosquito populations has never been found. A possible explanation is that mosquitoes use decreasing air pressure as an indicator for adverse weather conditions and modify their behaviour (e.g. mating) to reduce the risk of injury or death. Such changes in behaviour were previously reported for *Drosophila melanogaster* (Austin et al., 2014) as well as for beetles, moths, and aphids (Pellegrino et al., 2013).

Our results on the dew point suggest that evaporation and condensation of water are causal drivers of the mosquito dynamics. Evaporation reduces water level and increases salinity in breeding containers, whereas condensation has the opposite effect. A previous study showed a positive correlation between the dew point and the percentage of containers holding water versus number of *Ae. aegypti* larvae (Favier et al., 2006).

### Temporal scale dependence of mosquito-climate interactions

The mosquito-climate interactions exhibited temporal scale dependence. This may be associated with the mosquito life cycle that consists of 5-10 days larval development including pupation, and a total life span of up to four weeks. Our causality analysis revealed short time-lag effects of 0-9 days for temperature, precipitation, dew point, and mean tide level, which may reflect a causal effect of the climate variable on adult mortality, larval developmental time or mortality at late larval stages. Intermediate time-lag effects were found for temperature, precipitation, mean tide level, and air pressure, which may reflect an effect on larval mortality, larval competition or larval developmental time. Long time-lag effects were found for temperature, dew point, and mean tide level, which may indicate an effect of the climate variables on female fecundity or host availability (e.g., human, bird, rat, dog). Very long time-lag effects of 30-39 days were found for dew point, mean tide level, and air pressure, which may reflect an effect on the availability of breeding habitats such as broken coconuts or crab holes. We remark that the causal effect may be direct, e.g., temperature may directly affect larval survival due to stress, or indirect, e.g., temperature may alter behaviour of larval predators. In addition, indirect causality may involve human activity, e.g., temperature dependent behaviour of pesticide usage. Importantly, the same climate variable may affect the mosquito at different life stages with different time lags. This may even occur at the same population; e.g., S4 is affected by temperature with delays of 8 days and 20-23 days.

### Spatial variation in mosquito-climate interactions

The causality network displays strong spatial scale dependence with respect to mosquito-environment interactions. Causal interactions were only detected at the smallest spatial scale of a single sampling site, but neither at the level of a motu nor at the whole meta-population. Similar patterns were previously reported for the dynamics of measles, which showed signatures of nonlinearity and predictability only at the city-by-city scale, but not at the larger country-wide scale (Sugihara et al. 1992). In both cases, signatures for nonlinearity are lost by summation of subpopulation dynamics. This may seem counter-intuitive, but is actually to be expected for a nonlinear dynamical system, in which subpopulations are affected by different forcing variables or the forcing is non-synchronized (Sugihara and May 1990), as was found in this study (Fig. 1).

The causality analysis revealed substantial heterogeneity in terms of causal environmental variables and time-lagged effects between neighbouring populations even on the same motu (Fig. 1). How are such differences at a very small spatial scale determined? A possible explanation is that populations in close proximity respond differently to the climate even if the distance is less then 100 m. When interpreting the results, however, it should be kept in mind that the climate data were not directly collected at the mosquito sampling points (Methods). Climate data with spatial resolution are necessary to evaluate whether *Ae. polynesiensis* is really affected by its microhabitat.

An interesting finding was that mosquito-environment interactions are more complex on Toamaro and Tiano than on Horea (Fig. 1). This difference in complexity may, to some extent, be explained by differences in mosquito body size. In a previous study on *Ae. polynesiensis* using the same mosquito samples, Mercer et al. (2012) reported that female body size is significantly smaller on Toamaro motu than on Horea. The authors explain this finding by higher larval densities and longer larval developmental times on Toamaro (Mercer et al. 2012). We hypothesize that either (i) the longer developmental times and the smaller adult body sizes make mosquitoes on Toamaro more sensitive to environmental fluctuations, and that this causes the complex mosquito-climate interactions found for this motu or (ii) the climatic variables influence the larval stage which leads to longer development time, smaller adult body size, and the complex dynamics. The hypotheses could be tested using longitudinal data that include both adult and larval stages.

While the three motus were chosen as study sites because they have similar climate, they actually differ with respect to vegetation and human impacts. In Tiano, pest control is intense and canopy cover and mosquito abundances are low; in Horea, canopy cover is high, pest control happens on irregular basis and mosquito abundances are intermediate; in Toamaro, canopy cover is high, pest control is mostly absent, and mosquito abundance is high. Among the three motus, Toamaro suffered the least human effects, while the mosquito population on this motu are characterized by the most complex interactions between the climate variables and mosquito abundances (Fig. 1). This suggests that pest control strategies may not only reduce mosquito abundance but also alter the complexity of mosquito-climate interdependencies. This may be a relevant aspect in pest control.

### Causality between mosquito populations

An interesting finding was the detection of causality between subpopulations (Fig. 1). There is a clear spatial pattern with five out of six links connecting neighbouring populations, and only one link connecting two motus. A possible explanation is that female mosquitoes disperse by prevailing winds.

Our results demonstrate that CCM allows identifying interactions within a meta-population. Meta-population dynamics and structure are important determinants of ecological and evolutionary processes (Hanski and Gilpin, 1997; Hanski, 1998). Meta-population dynamics and structure were typically studied by directly measuring migration and dispersal, e.g., by mark-release-recapture experiments (Guerra et al 2014), or indirectly reconstructed by estimating gene flow between populations, e.g. using microsatellite data (Brelsfoard and Dobson, 2012). Here for the first time, we demonstrated that CCM may provide an alternative solution, which may be of wide applicability. For example, one may study how climate variables, community composition, and population density causally affect the migration behaviour of a focal species. This analysis provides information of the interconnectedness in a meta-population. CCM and scenario exploration allow further investigating whether the meta-population structure itself is context-dependent and how changes of climate variables alter the interconnectedness in the meta-population. Such research questions are difficult to study with the existing methods.

### Scenario explorations

We used the scenario exploration technique (Deyle et al., 2013) to predict how environmental change affects mosquito abundance. Our analysis revealed strong state dependent responses (Fig. 2, supplementary Fig. S7). This result is expected given the various empirical evidences for complex and nonlinear mosquito-environment interactions. Here, we tailored scenario exploration to the specific needs of disease vector control, and investigated how environmental change affects mosquito outbreaks. This analysis suggests that climate change may substantially alter the number and strength of outbreaks, but that the outcome critically depends on the local environmental conditions and may differ even between populations in short distance of less than 200 m (Fig. 3).

We propose using scenario exploration as a new framework to predict environmental change effects on mosquito abundances and disease outbreaks. The main difference to previous forecasting methods is that realistic, nonlinear mosquito-climate interdependencies are fully acknowledged. We believe that this offers new opportunities for mosquito forecasting. As the skill of EDM forecasting depends crucially on the quality of data, there is a critical need for long-term mosquito monitoring. We suggest to collect time series data of mosquito abundances on a regular base of 1-2 weeks, and combine them with climate information from data logger and case reports of transmitted diseases at local hospitals. Scenario exploration may then be used to (1) predict when outbreaks of mosquitoes or mosquito-borne diseases happen, and to (2) simulate different pest control strategies and explore how they depend on climate conditions. In that sense, we hope that this study sets a first step towards a new direction of pest control and disease prevention strategies.

In conclusion, we deciphered the causality network of a mosquito meta-population with climate variables (temperature, precipitation, dew point, air pressure, mean tide level) using EDM (Fig. 1) that was otherwise difficult to discern using linear methods. The findings that the responses of subpopulations in close proximity (100-500 m) differed substantially with respect to their causal environmental drivers and that no causal climate variables were found for the motu scale and whole meta-population scale highlights the importance of spatial scale dependency in complex systems. Moreover, we found strong temporal scale dependence of the causality with time lags ranging from zero up to five weeks, suggesting that climate acts on the different life stages of the mosquito and that the interdependencies of mosquito-climate interactions have a high degree of complexity. More importantly, albeit with the complexity, we presented a new framework based on EDM to predict how environmental change may affect the number and strength of mosquito outbreaks. These results bear profound management implications for disease vector ecology and pest control.

## Methods

### Mosquito field site

Adult *Aedes polynesiensis* mosquitoes were collected on three neighboring motu islands, Horea, Tiano, and Toamaro, on the western coast of Raiatea (Mercer et al., 2012, see also figure 1). Mosquito data are shown in figure S1.

The linear distance between the shoreline of Raiatea and the motus is approximately 500-700 m. Horea, the smallest among the three, is a publically held circular island of an area of 1.41 ha, with vegetation consisting of 60-80% canopy cover. This motu is semi permanently inhabited, and insecticides and wood smoke are used for mosquito control. Mosquito larvae on this motu develop in artificial containers, coconuts, spathes, phytotemata, and burrows made by the terrestrial crab *Cardisoma carnifex*. Blood meals of mosquitos are from humans, dogs, rats, and birds including chicken.

Tiano, the largest among the three, is privately owned and has a size of 6.34 ha. The vegetation has an open canopy in the center (<20% cover) and 80-90% canopy around the edge. Mosquitoes are controlled by pesticide usage and breeding site reduction. Mosquito larvae on this motu develop in cryptic tree holes, crab burrows, and a pool that is temporarily filled with water. Blood meals are from dogs, rats, birds, and humans.

Toamaro is a privately owned motu with an area of 4.06 ha. The vegetation consists of 70-95% canopy and has greater stratification with herbaceous and shrub layers than Horea and Tiano. This island is occasionally managed by vegetation clearing. Insecticides are used for mosquito control during periods of coconut collection. Mosquito larvae on this motu develop in artificial containers, broken coconuts, phytotemata, and crab burrows. Blood meals are from rats, birds, and occasionally dogs, cats, and human.

### Time series data

*Ae. polynesiensis* population data were previously presented by Mercer et al. (2012). Adult mosquitoes were collected on three neighboring motu islands, Horea, Tiano, and Toamaro, on the western coast of Raiatea (Society Archipelago, French Polynesia; 16°49.4’S, 159°29.2’W) from 1 February 2008 to 3 December 2009 (Fig. 1). On each motu, there were three sampling sites, and mosquitoes were collected at a 14-day interval using Biogent Sentinel traps (data shown in supplementary Fig. S1). Sampling sites are numbered serially from north to south and denoted by S1 to S9. All collected mosquitoes were species identified, sex determined, and counted for each time interval. In the present study, we focused on female *Ae. polynesiensis*, which comprised about 99% of the caught mosquitoes. For further details see Mercer et al. (2012).

Climate data were downloaded from the *National Oceanic and Atmospheric Administration* (http://www.noaa.gov/).From Bora Bora airport (16°26.4’S, 151°45.1’W; approx. 50km distance to the sampling sites at Raiatea), we obtained daily records of mean temperature (*T*), dew point (*DEW*), precipitation (*PREC*), and air pressure (mean atmospheric sea level pressure, *SLP*). From Pago Pago (American Samoa; 14°16.6’S, 170°41.3’W), we obtained hourly tide data, from which we calculated the daily average mean tide levels (*MTL*). Pago Pago has a distance of approximately 2000 km to Raiatea and was the nearest weather station providing tide data. Note that Raiatea and Pago Pago have similar latitude. All climate data were recorded daily for the period from 7 December 2007 to 3 December 2009 (supplementary Fig. S2). This period is eight weeks longer than mosquito sampling and thus allows investigating possible time-lag effects of climate variables.

### Data analysis

#### Convergent cross mapping

Causal effects between mosquito populations, and from environmental variables to mosquito populations were tested using *convergent cross mapping* (CCM) (Sugihara et al., 2012). CCM tests whether two variables, each given as a time series, are coupled in the same dynamical system. Variable *X* is said to be causally affected by *Y* if the underlying dynamic of *X* is a function of *Y*. This method, based on Takens’ theory for dynamical systems (Takens, 1981), tests for causation by measuring the extent to which the causal variable has left an imprint in the time series of the affected (Sugihara et al 2012). The method is rooted in state space reconstruction of shadow manifold by lagged coordinate embedding, and the essential ideas are summarized in the following animations: tinyurl.com/EDM-intro. In this study, the embedding dimension (*E*_*b*_) for each causal link (e.g. from *Y*(*t*) to *X*(*t*)) in CCM analysis was determined by testing values of *E* from 2 to 10 dimensions that optimizes the cross-mapping *ρ* in which *X*(*t*) is used to predict *Y*(*t*).

The causation, *Y* causes *X*, can be confirmed if and only if SSR_*X*(*t*)_ cross-mapping SSR_*Y*(*t*)_ converges, meaning that the cross-mapping skill *ρ*(*L*) improves with increasing library lengths (*L*) of *X* (Sugihara et al 2012). The direction of cross-mapping (*X* maps *Y*) is opposite to the direction of causation (*Y* causes *X*) because only *Y* can leave footprints on *X* and makes the backward cross-mapping from *X* possible. Note that in this study, we also explored the causations with lag response (van Nes et al., 2015). Specifically, for investigating effects of climate variables on mosquitos, we tested for time lags ranging from *d* = 0 to 42 days, while 14-day arithmetic means of the climate variables were used such that the climate time series match the mosquito-sampling interval. To investigate causal effect among mosquito populations, we conduced CCM between paired populations. Potential lag effects were evaluated for *d* = 0, 14 and 28 days.

To determine the convergence in cross-mapping, three statistical criteria were applied. First, Kendall’s *τ* test was used to examine whether the cross-map skill *ρ* increases monotonously as a function of library size *L*. Results are considered as significant if *τ* > 0 and *P* < 0.01. Second, Fisher’s Z test was used to examine whether the predictability under maximum library size, *ρ(L*_*max*_), is significantly higher than the predictability under minimum library size, *ρ(L*_*min*_). The threshold was set to *P* < 0.05. Third, we tested whether *ρ(L*_*max*_) of the observed time series is significant different from a null model, using the surrogate time series test. 100 surrogates were generated, following Ebisuzaki (1997), by randomly reshuffling phases, while preserving the mean, variation and power spectrum. The results are deemed significant if *ρ(L*_*max*_) of the original data is larger than the 95% upper bound of the surrogates (*P* < 0.05) (Deyle et al., 2013; van Nes et al., 2015). To account for multiple testing, however, we reduced the level of significance to *P* < 0.01 when testing for causal climate effects.

#### Scenario exploration

To evaluate how change in a climate variable (the causal variable identified by CCM) affect mosquito abundances of a given subpopulation, we used multivariate simplex projection following (Deyle et al., 2013). Specifically, we conducted scenario explorations with an extended shadow manifold that has *E*_*b*_+1 dimensions (Deyle et al., 2013; supplementary Fig. S6). The first *E*_*b*_ dimensions contain the time-delayed embeddings of the mosquito abundances, as described above. Then, to explore potential effects of changing environment, an additional coordinate axis of either the time series of a climate variable or another mosquito population was appended, resulting in *E*_*b*_+1 dimensions in state space. It is this extra variable that we aim to “manipulate” to achieve scenario exploration. Specifically, we purposely increase or decrease that forcing variable by multiples of its standard deviation and then forecast the resultant mosquito abundance one step (in our case, 14 days) into the future for every time point.

The scenario exploration approach allows us to investigate how change in a climate variable affects the number and strength of mosquito outbreaks for each mosquito time series. Here, mosquito outbreaks are defined as mosquito abundances that are five standard deviations larger than the median abundance. The outbreak strength is defined as the arithmetic mean of the five highest abundances. For each mosquito time series, we compared the number and strength of mosquito outbreaks as predicted under a “scenario” versus those in the original data. To quantify the effect of a scenario of changing a climate variable, we calculated the relative outbreak strength, which is defined as the arithmetic mean of the five highest population abundances under scenario exploration divided by that of the original data.

#### Further details of data analysis

Time series analysis was conducted using the rEDM package (version 0.2.4) of the programming language *R* (Ye at al., 2016). To minimize autocorrelation and heteroscedasticity, all time series were first differenced and normalized to zero mean and unit variance (Sugihara, 1994; Telschow et al., 2017). Due to the paucity of observation points, we performed leave-one-out cross-validation (Glaser et al., 2014).

## Authors contributions

Design of study: FG, CH, AT; data analysis: FG; interpretation of results: FG, JS, CH, AT; writing of manuscript: FG, JS, CH, AT. All authors gave final approval for publication.

## Additional information

The authors declare no competing financial interests.

## Acknowledgements

This work was supported by the Project-Based Personnel Exchange Program between the NSC and DAAD (101-2911-I-002-507; 54368760) awarded to CH and AT. AT was supported by the German Science Foundation (SPP 1399, TE 976/2-1) and the Volkswagen Foundation’s evolutionary biology initiative, CH by the National Center for Theoretical Sciences, Foundation for the Advancement of Outstanding Scholarship, and the Ministry of Science and Technology (Taiwan).

## References

1. Alencar J, de Mello CF, Guimarães AÉ, Gil-Santana HR, Silva Jdos S, Santos-Mallet JR, Gleiser RM (2015). Culicidae Community Composition and Temporal Dynamics in Guapiaçu Ecological Reserve, Cachoeiras de Macacu, Rio de Janeiro, Brazil. PLOS ONE v10(3): e0122268. doi: 10.1371/journal.pone.0122268

2. Austin CJ, Guglielmo CG, Moehring AJ (2014). A direct test of the effects of changing atmospheric pressure on the mating behavior of Drosophila melanogaster. Evol Ecol v28(3):535–544. doi: 10.1007/s10682-014-9689-8

3. Azil AH, Long SA, Ritchie SA, Williams CR (2010). The development of predictive tools for pre ‐emptive dengue vector control: a study of Aedes aegypti abundance and meteorological variables in North Queensland, Australia. Tropical Medicine & International Health v15(10):1190–1197. doi: 10.1111/j.1365-3156.2010.02592.x

4. Barrera R, Amador M, Clark GG (2006). Ecological factors influencing Aedes aegypti (Diptera: Culicidae) productivity in artificial containers in Salinas, Puerto Rico. J Med Entomol v43(3):484–492. doi: 10.1603/0022-2585(2006)43[484:EFIAAD]2.0.CO;2

5. Barrera R, Amador M, MacKay AJ (2011). Population dynamics of Aedes aegypti and dengue as influenced by weather and human behavior in San Juan, Puerto Rico. PLoS Negl Trop Dis v5(12): e1378. doi: 10.1371/journal.pntd.0001378

6. Bashar K and Tuno N (2014). Seasonal abundance of Anopheles mosquitoes and their association with meteorological factors and malaria incidence in Bangladesh. Parasit Vectors v7(422). doi: 10.1186/1756-3305-7-442

7. Brelsfoard CL and Dobson SL (2012). Population genetic structure of Aedes polynesiensis in the Society Islands of French Polynesia: implications for control using a Wolbachia-based autocidal strategy. Parasit Vectors v5(80). doi: 10.1186/1756-3305-5-80

8. Carrington LB, Armijos MV, Lambrechts L, Scott TW (2013). Fluctuations at a low mean temperature accelerate dengue virus transmission by Aedes aegypti. PLoS neglected tropical diseases, 7(4), e2190.

9. Chang YM (2010). Distribution of Two Aedes Mosquito Disease Vectors on Moorea, French Polynesia: A Study of the Effects of Environmental Factors and Changes in Larval Habitat. Berkely University: Biology and Geomorphology of Tropical Islands, Student Papers. Online publication.

10. Chang CW, Ushio M, Hsieh CH (2017). Empirical Dynamic Modeling for beginners. Ecological Research. 32: 785–796

11. Chaves LF, Morrison AC, Kitron UD, Scott TW (2012). Nonlinear impacts of climatic variability on the density ‐dependent regulation of an insect vector of disease. Global Change Biology v18(2):457–468. doi: 10.1111/j.1365-2486.2011.02522.x

12. Couret J, Dotson E, Benedict MQ (2014). Temperature, Larval Diet, and Density Effects on Development Rate and Survival of Aedes aegypti (Diptera: Culicidae). PLOS ONE. doi: 10.1371/journal.pone.0087468

13. Deyle ER, Fogarty M, Hsieh CH, Kaufman L, MacCall AD, Munch SB, Perretti CT, Ye H, Sugihara G (2013). Predicting climate effects on Pacific sardine. Proc Natl Acad Sci U S A v110(16):6430–5. doi: 10.1073/pnas.1215506110

14. Deyle ER, Maher MC, Hernandez RD, Basu S, Sugihara G (2016). Global environmental drivers of influenza. Proc Natl Acad Sci U S A v113(46):13081–13086. doi: 10.1073/pnas.1607747113

15. Dye C (1984b). Models for the population dynamics of the yellow fever mosquito, Ae. aegypti. J Anim Ecol. v53:247–268. doi: 10.2307/4355

16. Duncombe J, Clements A, Davis J, Hu W, Weinstein P, Ritchie S (2013). Spatiotemporal patterns of Aedes aegypti populations in Cairns, Australia: assessing drivers of dengue transmission. Trop Med Int Health. v18(7):839–849. doi: 10.1111/tmi.12115

17. Ebisuzaki W (1997). A method to estimate the statistical significance of a correlation when the data are serially correlated. Journal of Climate v10(9):2147–2153. doi: 10.1175/1520-0442(1997)010<2147:AMTETS>2.0.CO;2

18. Favier C, Degallier N, Ribeiro Vilarinhos PDT, de Carvalho MDL, Cavalcanti Yoshizawa MA, Knox MB (2006). Effects of climate and different management strategies on Aedes aegypti breeding sites: a longitudinal survey in Brasilia (DF, Brazil). Tropical Medicine & International Health v11(7):1104–1118. doi: 10.1111/1365-3156.2006.01653.x

19. Focks DA, Haile DG, Daniels E, Mount GA (1993). Dynamic life table model for Aedes aegypti (Diptera: Culicidae): analysis of the literature and model development. J Med Entomol v30(6):1003–1017. doi: 10.1093/jmedent/30.6.1003

20. Galun R and Fraenkel G (1961). The effect of low atmospheric pressure on adult Aedes aegypti and on housefly pupae. Journal of Insect Physiology v7(3-4):161–176. doi: 10.1016/0022-1910(61)90069-5

21. GBD 2015 Mortality and Causes of Death Collaborators (2016). Global, regional, and national life expectancy, all-cause mortality, and cause-specific mortality for 249 causes of death, 1980 – 2015: a systematic analysis for the Global Burden of Disease Study 2015. Lancet v388(10053):1459–1544. doi: 10.1016/S0140-6736(16)31012-1

22. Glaser SM, Fogarty MJ, Liu H, Altman I, Hsieh C-H, Kaufman L, MacCall AD, Rosenberg AA, Ye H, Sugihara G (2014). Complex dynamics may limit prediction in marine fisheries. Fish Fish 15(3):616–633. doi: 10.1111/faf.12037

23. Guerra CA, Reiner RC Jr., Perkins TA, Lindsay SW, Midega JT, Brady OJ, et al. (2014). A global assembly of adult female mosquito mark-release-recapture data to inform the control of mosquito-borne pathogens. Parasit Vectors v7(1):276. doi: 10.1186/1756-3305-7-276

24. Hanski I and Gilpin ME (1997). Metapopulation Biology: Ecology, Genetics, and Evolution. ISBN: 978-0-12-323445-2

25. Hanski I (1998). Metapopulation dynamics. Nature 396(6706):41–49. doi: 10.1038/23876

26. Hsieh CH, Glaser SM, Lucas AJ, Sugihara G (2005). Distinguishing random environmental fluctuations from ecological catastrophes for the North Pacific Ocean. Nature 435:336–340. doi:10.1038/nature03553

27. Ingram RL (1954). A Study of the Bionomics of Aedes (Stegomyia) polynesiensis Marks under Laboratory Conditions. Am.J.Hyg. 60:169–185.

28. Lambdin BH, Schmaedick MA, McClintock S, Roberts J, Gurr NE, Marcos K, Waller L, Burkot TR (2009). Dry Season Production of Filariasis and Dengue Vectors in American Samoa and Comparison with Wet Season Production. B Am J Trop Med Hyg v81(6): 1013–1019. doi: 10.4269/ajtmh.2009.09-0115

29. Lambrechts L, Paaijmans KP, Fansiri T, Carrington LB, Kramer LD, Thomas MB, Scott TW (2011). Impact of daily temperature fluctuations on dengue virus transmission by Aedes aegypti. Proc Natl Acad Sci U S A 108(18):7460–7465. doi: 0.1073/pnas.1101377108

30. Lau CL, Won KY, Lammie PJ, Graves PM (2016). Lymphatic Filariasis Elimination in American Samoa: Evaluation of Molecular Xenomonitoring as a Surveillance Tool in the Endgame. PLOS Neglected Tropical Diseases 10(11): e0005108. doi: 10.1371/journal.pntd.0005108

31. Legros M, Lloyd AL, Huang Y, Gould F (2009). Density-Dependent Intraspecific Competition in the Larval Stage of Aedes aegypti (Diptera: Culicidae): Revisiting the Current Paradigm. J Med Entomol v46(3):409–419. doi: 10.1603/033.046.0301

32. Mains JW, Brelsfoard CL, Crain PR, Huang Y, Dobson SL (2013). Population impacts of Wolbachia on Aedes albopictus. Ecological applications, 23(2), 493–501.

33. Mercer DR, Bossin H, Sang MC, O’Connor L, Dobson SL (2012). Monitoring temporal abundance and spatial distribution of Aedes polynesiensis using BG-Sentinel traps in neighboring habitats on Raiatea, Society Archipelago, French Polynesia. J Med Entomol v49(1):51–60. doi: 10.1603/ME11087

34. Pellegrino, AC Peñaflor MF, Nardi C, Bezner-Kerr W, Guglielmo CG, Bento JM, McNeil JN (2013). Weather forecasting by insects: modified sexual behaviour in response to atmospheric pressure changes. PLoS One 8(10):e75004. doi: 10.1371/journal.pone.0075004

35. Scott TW, Morrison AC, Lorenz LH, Clark GG, Strickman D, Kittayapong P, Zhou H, Edman JD (2000). Longitudinal studies of Aedes aegypti (Diptera: Culicidae) in Thailand and Puerto Rico: population dynamics. J Med Entomol v37(1):77–88. doi: 0.1603/0022-2585-37.1.77

36. Simões T C, Codeço CT, Nobre AA, and Eiras AE (2013). Modeling the Non-Stationary Climate Dependent Temporal Dynamics of Aedes aegypti. PLOS ONE 8:e64773. doi: 10.1371/journal.pone.0064773

37. Sugihara G, Grenfell B, May RM, Chesson P, Platt HM, Williamson M (1990). Distinguishing error from chaos in ecological time series. Phil. Trans. R. Soc. B 330(1257). doi: 10.1098/rstb.1990.0195

38. Sugihara G (1994). Nonlinear forecasting for the classification of natural time series. Phil. Trans. R. Soc. Lond. A 348:477–495. doi: 10.1098/rsta.1994.0106

39. Sugihara G, May R, Ye H, Hsieh CH, Deyle E, Fogarty M, Munch S (2012). Detecting causality in complex ecosystems. Science 338:496–500. doi: 10.1126/science.1227079

40. Takens F (1981). Detecting strange attractors in turbulence. In D. A. Rand and L.-S. Young. Dynamical Systems and Turbulence, Lecture Notes in Mathematics, vol. 898. Springer-Verlag. pp. 366–381.

41. Telschow A, Grziwotz F, Crain P, Miki T, Mains JW, Sugihara G, Dobson SL, Hsieh CH (2017). Infections of Wolbachia may destabilize mosquito population dynamics. Journal of Theoretical Biology 428: 95–105. doi.org/10.1016/j.jtbi.2017.05.016.

42. van Nes EH, Scheffer M, Brovkin V, Lenton TM, Ye H, Deyle E, Sugihara G (2015). Causal feedbacks in climate change. Nature Climate Change v5(5):445–448. doi: 10.1038/NCLIMATE2568

43. Ye H, Beamish RJ, Glaser SM, Grant SC, Hsieh CH, Richards LJ, Schnute JT, Sugihara G (2015). Equation-free mechanistic ecosystem forecasting using empirical dynamic modeling. v112(13):E1569–76.doi: 10.1073/pnas.1417063112

44. Ye H, Clark A, Deyle E, and Sugihara G (2016). rEDM: an R package for Empirical Dynamic Modeling and Convergent Cross-Mapping.

