## Supplementary Materials for "Identify and Predict Environmental Change Effects On Tiger Mosquitos, *Aedes Polynesiensis*"

S1 Supplementary figures

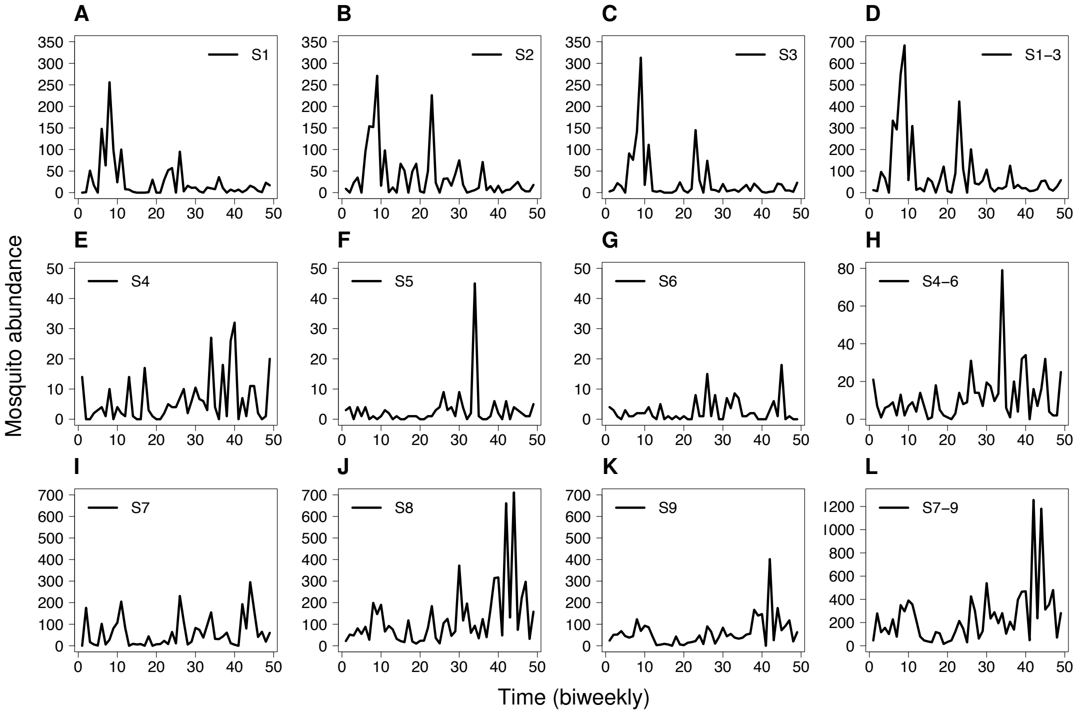


**Figure S1. Time series data of female *Ae. polynesiensis* mosquitoes.** Plots A-D show mosquito abundances at Horea; symbols S1 to S3 indicate the sampling sites, and S1-3 the total abundance of the motu. Plots E-H show mosquito abundances at Tiano; S4 to S6 indicate the sampling sites, and S4-6 the total abundance. Plots I-L show mosquito abundances at Toamaro; S7 to S9 indicate the sampling sites, and S7-9 the total abundance.

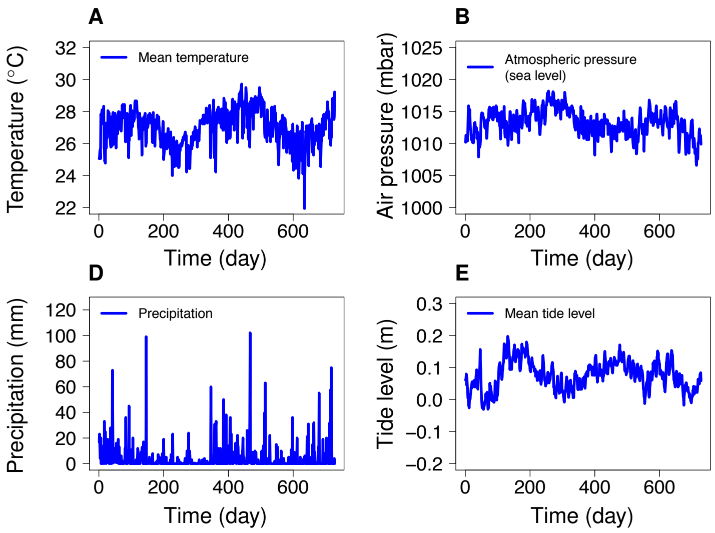


**Figure S2. Daily climate time series.** Further details on the data are given in the Methods.

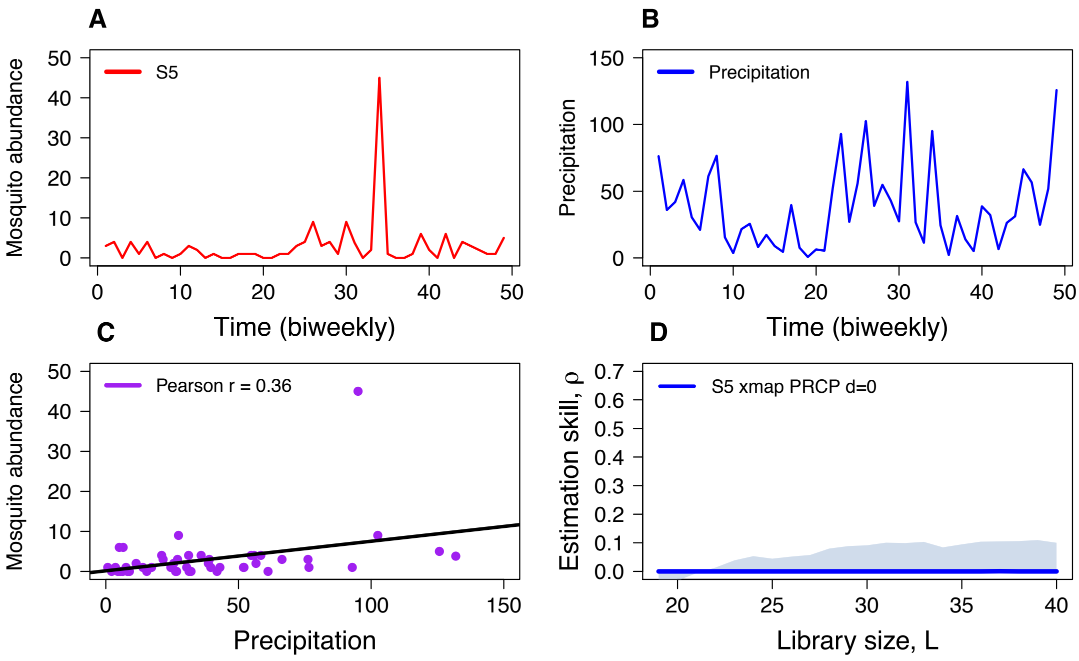


**Figure S3. Correlation does not imply causality.** *Panel A*: *Ae. polynesiensis* abundance at sampling site S5 as function of time. *Panel B:* 14-day averaged precipitation (PRCP) with time lag *d*=0 as functions of time. *Panel C:* Mosquito abundance at S5 as function of average precipitation. There is significant positive correlation between the two variables (Pearson r=0.36, P<0.05; Spearman ρ=0.38, P<0.01). *Panel D:* CCM test for causal effect of precipitation on mosquito abundance at S5 (S5 xmap PRCP). The blue line indicates the estimation skill as function of the library size. There is no detectable causal effect as the estimation skill does not converge and is within the 95 percentile of 100 randomly generated surrogates (Methods). The figure illustrates that correlation does not necessarily imply causation.

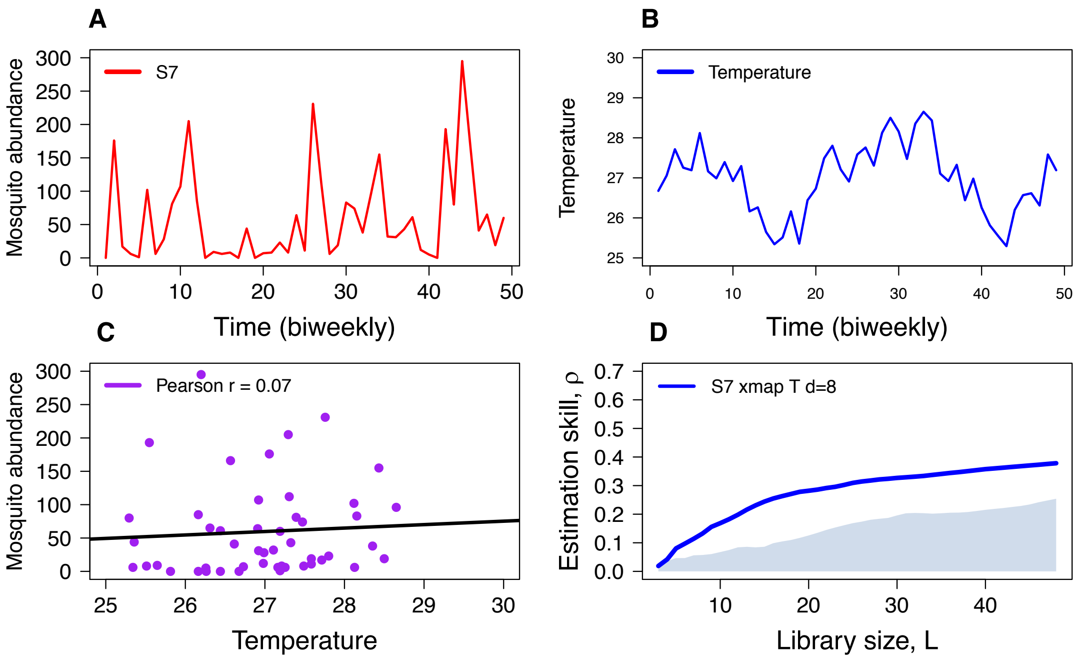


**Figure S4. Causality does not imply correlation.** *Panel A*: *Ae. polynesiensis* abundance at sampling site S7 as function of time. *Panel B:* 14-day averaged temperature (T) with time lag *d*=8 as functions of time. *Panel C:* Mosquito abundance at S7 as function of average temperature. There is no significant correlation between the two variables (Pearson r=0.07, P>0.05; Spearman ρ=0.19, P>0.05). *Panel D:* CCM test for causal effect of temperature on mosquito abundance at S7 (S7 xmap T). The blue line indicates the estimation skill as function of the library size. There is a significant causal effect because the estimation skill converges and is not within the 95 percentile of 100 randomly generated surrogates (Methods). The figure illustrates that causation does not necessarily imply correlation.

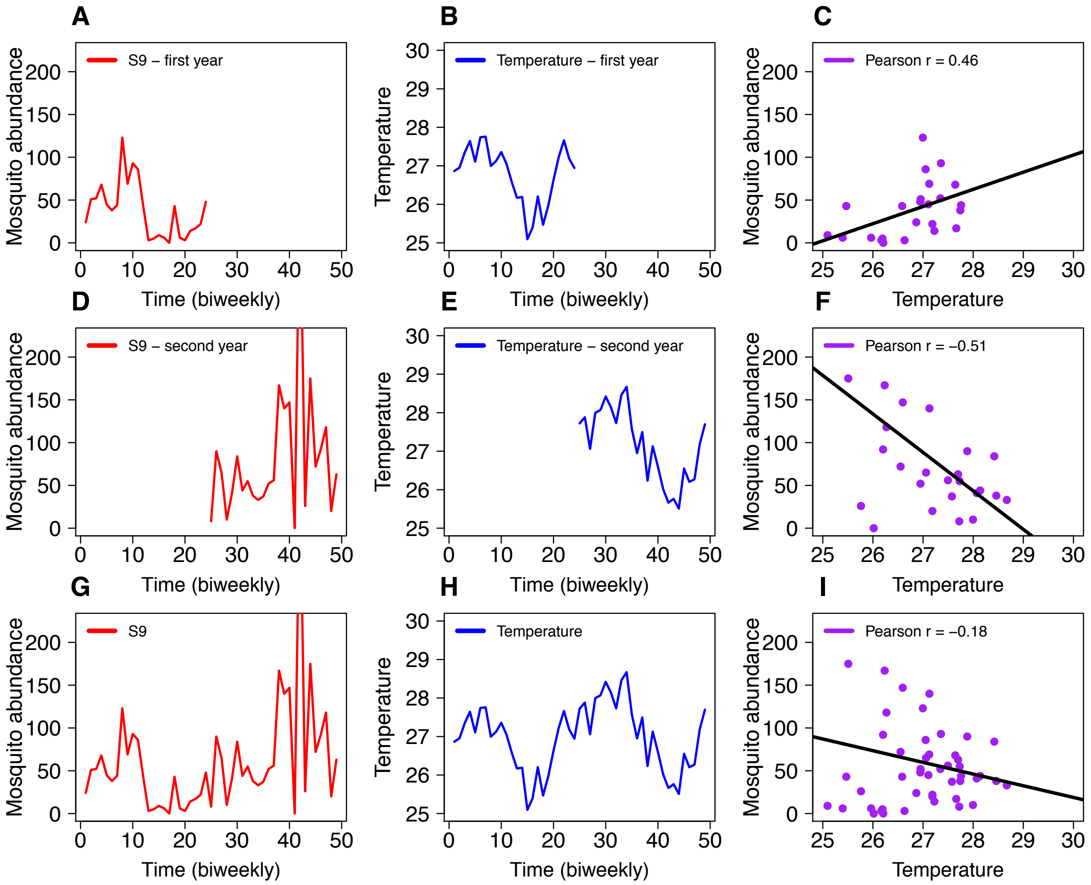


**Figure S5. Mirage correlation.** Shown are mosquito abundances of sampling site S9 (A, D, G), the average temperature with a time lag of *d*=14 days (B, E, H), and correlation plots of temperature and mosquito abundance (C, F, I). In the first year (2008), mosquito abundance and temperature are significantly positive correlated (Pearson r=0.46, P<0.05; Spearman ρ=0.52, P<0.01). In the second year (2009), mosquito abundance and temperature are significantly negative correlated (Pearson r=-0.51, P<0.01; Spearman ρ=-0.42, P<0.05). However, there are no significant correlations when the whole two-year period is analyzed (Pearson r=-0.18, P>0.05; Spearman ρ=0.07, P>0.05).

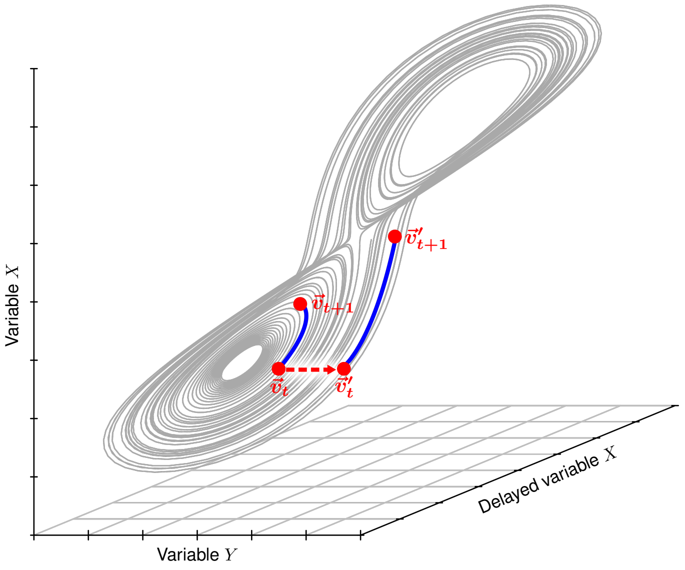


**Figure S6.** Schematic illustrating the method of scenario exploration, using the Lorenz (Lorenz, 1963) butterfly attractor as an example. For this well studied system, the embedding dimension is 3. To illustrate, we used *Y*(t), *X*(t) and *X*(t-1) for the 3 coordinates in state space reconstruction. The gray line represents the trajectory of the system states and characterizes the attractor. The red dots are arbitrarily chosen to indicate specific states for demonstration. Based on Takens' Theorem, knowledge of the attractor allows predicting how the system changes with time. For example, consider $\vec{v}_t = (x_t, x_{t-1}, y_t)$, i.e. the state of the system at time point *t*. Following the trajectory starting at $\vec{v}_t$ one time step into the future (indicated by the blue line) allows predicting the system state $\vec{v}_{t+1}$ at time *t*+1. To carry out scenario exploration, let's consider again state $\vec{v}_t$. In a first step, variable *Y* is perturbed by the amount *c* (indicated by the broken red arrow). This perturbation shifts vector $\vec{v}_t = (x_t, x_{t-1}, y_t)$ to vector $\vec{v}'_t = (x_t, x_{t-1}, y_t + c)$. As a consequence, the one-step forward prediction for vector $\vec{v}'_t$ becomes $\vec{v}'_{t+1}$. Comparison of $\vec{v}'_{t+1}$ with $\vec{v}_{t+1}$ allows evaluating how the system responds to change in variable *Y* at this specific time point (specific system state). Repeating the procedure for all time points (all states) allows evaluating more generally how the change of one variable affects the prediction. This basic idea allows evaluating how change in one variable affects the prediction of the system state one step into future. More generally, scenario exploration allows evaluating how change in causal variable affects the one-step forward prediction of the effect variable. The Lorenz system is described by three coupled differential equations, $\frac{dX}{dt} = s(Y - X)$, $\frac{dY}{dt} = X(r - Z) - Y$, and $\frac{dZ}{dt} = XY - bZ$. The system was numerically integrated for parameters *r*=28, *s*=10, and *b*=2.67 using the Runge-Kutta algorithm with step size of 0.01.

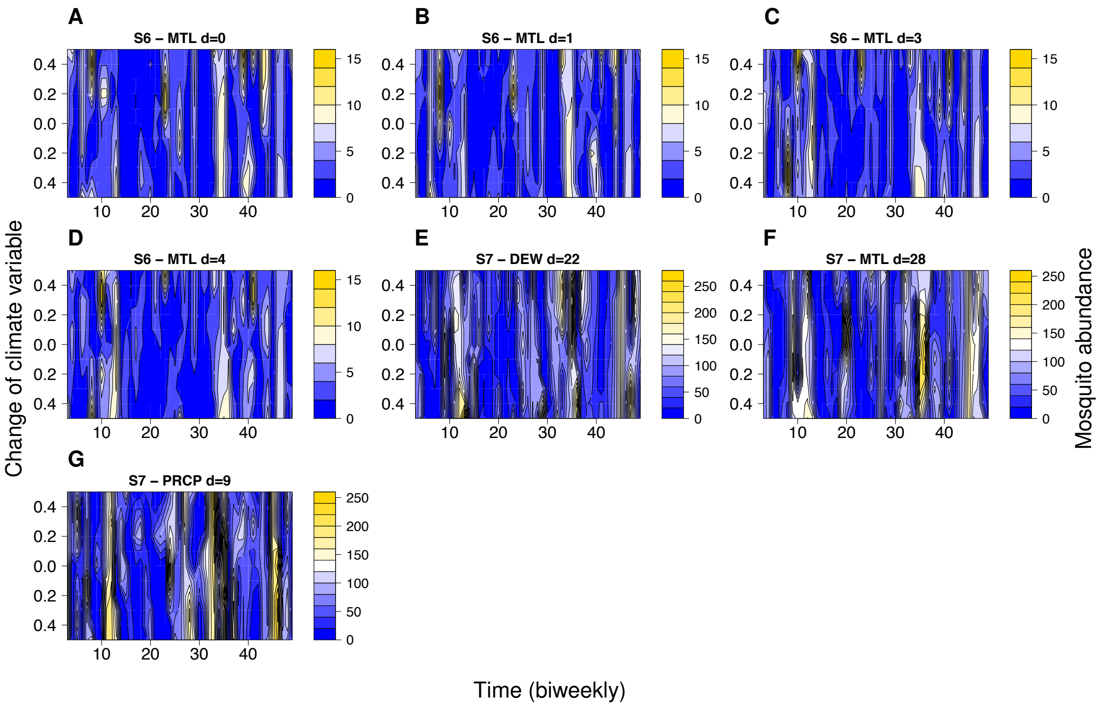


**Figure S7** Results of scenario exploration for mosquito dynamics. For exploring scenario, we purposely increase or decrease a forcing variable by multiples of its standard deviation ($\sigma$) and then forecast the resultant mosquito abundance one step into the future for every time point. Color indicates the predicted abundances of *Ae. polynesiensis* as a function of time (x-axis) and environmental change (y-axis). The y-axis ranges in $\pm 0.5\sigma$ of the climate variable. Note that the figure does not show scenario exploration for all 40 causality links shown in table S2, but only for the subset of seven links that were used in figure 3.

### S2 Supplementary tables

| site | $\rho_b$ | $E_b$ |
|---|---|---|
| S1 | 0.542 | 1 |
| S2 | 0.279 | 3 |
| S3 | 0.365 | 1 |
| S1-3 | -0.018 | 3 |
| S4 | 0.406 | 2 |
| S5 | 0.219 | 9 |
| S6 | 0.549 | 1 |
| S4-6 | 0.406 | 4 |
| S7 | 0.388 | 1 |
| S8 | 0.582 | 1 |
| S9 | 0.339 | 2 |
| S7-9 | 0.425 | 1 |
| S1-9 | 0.331 | 9 |

**Table S1.** Best embedding dimensions of the mosquito dynamics. Shown is the highest predictability, $\rho_b$, and the associated best embedding dimension, $E_b$, of each time series. Predictability is calculated as Pearson correlation between the observed data and the predictions (see Methods for details). The nine sampling points are denoted by S1 to S9, the Horea population by S1-3, the Tiano population by S4-6, the Toamaro population S7-9, and the total population of all three motu islands by S1-9.

| site | climate variable | $d$ | $\rho(L_{min})$ | $\rho(L_{max})$ | $P_{Fisher}$ |
|---|---|---|---|---|---|
| S1 | *SLP* | 14 | 0.007 | 0.378 | 0.032 |
| S1 | *SLP* | 15 | 0.03 | 0.397 | 0.032 |
| S1 | *SLP* | 17 | 0.003 | 0.381 | 0.03 |
| S3 | *MTL* | 3 | 0 | 0.382 | 0.028 |
| S3 | *MTL* | 4 | 0 | 0.411 | 0.019 |
| S3 | *MTL* | 5 | 0 | 0.438 | 0.013 |
| S3 | *MTL* | 6 | 0 | 0.461 | 0.009 |
| S3 | *MTL* | 7 | 0 | 0.454 | 0.01 |
| S3 | *MTL* | 8 | 0 | 0.432 | 0.014 |
| S3 | *MTL* | 9 | 0 | 0.4 | 0.022 |
| S4 | *T* | 13 | 0.011 | 0.364 | 0.041 |
| S4 | *T* | 20 | 0.028 | 0.442 | 0.018 |
| S4 | *T* | 21 | 0.024 | 0.487 | 0.009 |
| S4 | *T* | 22 | 0.007 | 0.481 | 0.008 |
| S4 | *T* | 23 | 0 | 0.39 | 0.027 |
| S4 | *DEW* | 33 | 0 | 0.365 | 0.036 |
| S4 | *DEW* | 34 | 0 | 0.407 | 0.021 |
| S4 | *DEW* | 35 | 0 | 0.405 | 0.022 |
| S4 | *MTL* | 21 | 0.135 | 0.465 | 0.042 |
| S4 | *MTL* | 22 | 0.146 | 0.483 | 0.037 |
| S4 | *MTL* | 23 | 0.14 | 0.459 | 0.048 |
| S6 | *MTL* | 0 | 0 | 0.377 | 0.03 |
| S6 | *MTL* | 1 | 0 | 0.364 | 0.035 |
| S6 | *MTL* | 2 | 0 | 0.357 | 0.038 |
| S6 | *MTL* | 3 | 0 | 0.364 | 0.035 |
| S6 | *MTL* | 4 | 0 | 0.354 | 0.039 |

| | | | | | |
|---|---|---|---|---|---|
| S7 | *T* | 8 | 0.019 | 0.378 | 0.036 |
| S7 | *DEW* | 8 | 0.006 | 0.349 | 0.044 |
| S7 | *DEW* | 22 | 0.045 | 0.395 | 0.039 |
| S7 | *DEW* | 23 | 0.02 | 0.37 | 0.04 |
| S7 | *PRCP* | 9 | 0.029 | 0.426 | 0.022 |
| S7 | *MTL* | 27 | 0.045 | 0.422 | 0.027 |
| S7 | *MTL* | 28 | 0.014 | 0.423 | 0.019 |
| S7 | *MTL* | 29 | 0 | 0.406 | 0.02 |
| S7 | *MTL* | 30 | 0 | 0.371 | 0.032 |
| S8 | *SLP* | 34 | 0 | 0.384 | 0.027 |
| S8 | *SLP* | 35 | 0 | 0.423 | 0.016 |
| S8 | *SLP* | 36 | 0 | 0.422 | 0.016 |
| S8 | *SLP* | 37 | 0 | 0.409 | 0.02 |
| S8 | *SLP* | 38 | 0 | 0.381 | 0.028 |

**Table S2. Causal effects of climate variables on mosquito dynamics**. Shown are the significant convergent cross mapping (CCM) from the mosquito population to the climate variables, as presented in figure 1. The time lag of the causal effect is denoted by *d*. Causality is identified by an increase of the CCM skill, i.e. $\rho(L_{max})>\rho(L_{min})$. The last column shows the P-values for the Fisher-z test ($P_{Fisher}$) for convergence of cross mapping.

| Site 1 | Site 2 | $d$ | $\rho(L_{min})$ | $\rho(L_{max})$ | $P_{Fisher}$ |
|---|---|---|---|---|---|
| S3 | S1 | 0 | 0.19 | 0.554 | 0.02 |
| S3 | S2 | 0 | 0.244 | 0.755 | 0.001 |
| S7 | S2 | 14 | 0 | 0.383 | 0.033 |
| S8 | S7 | 0 | 0.006 | 0.475 | 0.008 |
| S8 | S9 | 0 | 0.173 | 0.588 | 0.009 |
| S9 | S8 | 0 | 0.166 | 0.619 | 0.005 |

**Table S3. Causal interactions between mosquito subpopulations.** Shown are significant convergence cross mapping (CCM) results between mosquito abundances from different sampling points, as presented in figure 1. The first two columns indicate the effect and causal population, respectively. The third column shows the time lag of the causal interaction. Causality is identified by an increase of the CCM skill, i.e. $\rho(L_{max})>\rho(L_{min})$. The right column shows the P-values for the Fisher-z test ($P_{Fisher}$) for convergence of cross mapping.
